# Natural variation in expression of the mitochondrial flavoprotein WAH-1 alters response to cyanide in *C. elegans*

**DOI:** 10.1101/2023.03.03.531061

**Authors:** Maria P. Mercado, June H. Tan, Michael R. Schertzberg, Andrew G. Fraser

**Affiliations:** Department of Molecular Genetics, University of Toronto, Toronto, Canada; The Terrence Donnelly Centre for Cellular & Biomolecular Research, University of Toronto, Toronto, Canada

## Abstract

*C. elegans* is a free-living nematode that must adapt to a wide range of environments including both aerobic and anaerobic conditions. To survive in low oxygen, *C. elegans* can use an unusual form of anaerobic respiration that relies on rhodoquinone (RQ) as an alternative electron carrier. Parasitic nematodes like hookworm and whipworm also require rhodoquinone-dependent metabolism (RQDM) to survive in the highly anaerobic conditions in the human gut. Understanding how RQDM is regulated in *C. elegans* may thus identify new ways to combat these closely-related major human pathogens. We previously established a simple movement-based assay for RQDM in *C. elegans*. In this study, we tested a panel of wild-type isolates of *C. elegans* in our RQDM assay and find substantial variation in their ability to use RQDM. We carried out a genome-wide association study (GWAS) to identify loci that affect RQDM — this identified a single major QTL on the right arm of Chromosome III. We used RNAi to test almost all genes within the QTL region for involvement in RQDM and found one gene, *wah-1*, that strongly modulates RQDM-dependent recovery in *C. elegans*. WAH-1 is a mitochondrial flavoprotein that affects the electron transport chain, consistent with a role in RQDM. We show that *wah-1* expression varies between isolates due to major changes in *wah-1* transcript structures and this correlates tightly with variation in RQDM. Finally, we show that there is similar complexity to *wah-1* transcription in parasitic nematodes and that *wah-1* transcript structures change as parasites shift from aerobic to anaerobic, RQ- requiring metabolism. We thus conclude that reduced *wah-1* expression correlates with increased ability to survive in conditions where RQDM is essential.

## Introduction

Animals must be able to adapt to different environmental conditions. One critical challenge is the ability to generate energy in conditions where there is insufficient oxygen to support aerobic respiration using the mitochondrial electron transport chain. While most animals only experience hypoxia transiently, some animals live for extended periods in low oxygen conditions and have evolved various metabolic solutions to compensate^1, 2^. For example, goldfish survive for weeks in highly anaerobic conditions in frozen ponds by fermentation, excreting ethanol as a fermentation product^3^. Naked mole rats have evolved a different solution and switch from glucose to fructose as a fuel for anaerobic glycolysis in low oxygen to avoid feedback inhibition^4^.

The metabolic solutions that different animals have evolved to survive for extended periods are not just esoteric anomalies, but can have critical importance for human health. Soil-transmitted helminths (STHs)—including roundworm, hookworm and whipworm—are major human pathogens, infecting over a billion people worldwide^5, 6^. These STHs can survive for many weeks in the highly anaerobic conditions found within the human gut, and do this by using a highly unusual form of anaerobic respiration^7–9^. In normoxia, STHs generate energy using standard aerobic respiration. Electrons enter the mitochondrial electron transport chain (ETC) either at Complex I or via quinone-coupled dehydrogenases like Complex II (succinate dehydrogenase) where they are transferred onto ubiquinone (UQ)^10^. Electrons then pass along the ETC from UQ to cytochrome c in Complex III before finally being transferred to oxygen as the terminal electron acceptor. As electrons move through the ETC, Complex I, III and IV pump protons into the inner membrane space, generating a proton motive force across the inner mitochondrial membrane which powers ATP synthesis at Complex V, the F0F1- ATPase.

In anaerobic conditions, electrons are unable to exit the ETC at Complex IV since there is insufficient oxygen to act as a terminal electron acceptor. Under these conditions, many animals — including humans — cannot use the ETC. STHs, however, have evolved a different solution — they rewire their ETC to allow them to use alternative electron acceptors^8, 11–13^. Instead of being transferred onto UQ, electrons that enter the ETC at Complex I are transferred onto rhodoquinone (RQ), a highly related quinone molecule^14–16^. RQ has a greater reducing power than UQ, and can thus drive quinone-coupled dehydrogenases (QDHs) in reverse to act as reductases (e.g. succinate dehydrogenase is driven to act as a fumarate reductase)^11, 17–18^. This QDH reversal allows electrons to exit the ETC onto a range of terminal electron acceptors: in this case, electrons are transferred from RQ to fumarate by driving Complex II in reverse, thereby generating succinate as a terminal product. Humans do not make or use RQ and therefore RQ synthesis and RQ-dependent metabolism (RQDM) are ideal targets for new classes of anthelmintics.

We previously showed that *C. elegans* makes RQ and can use RQDM to survive in anoxic conditions^19, 20^. We developed a simple movement-based assay for RQDM and used this assay to dissect the pathway for RQ synthesis^20, 21^. In this study, we use this assay to measure variation in RQDM in a panel of 48 *C. elegans* natural isolates from the *Caenorhabditis elegans* Natural Diversity Resource (CeNDR) collection^22^. The CeNDR platform serves a repository for wild isolates that have been collected from all over the world, complete with fully sequenced genomes and annotated SNPs, offering a reservoir of natural genetic variation that can be used in GWA studies to correlate genotypic variants to phenotypically relevant traits^22–25^.

We observed significant variation in RQDM in the panel of isolates. To identify the QTLs underlying the variation in RQDM, we carried out a genome-wide association study (GWAS) and identified a major QTL affecting RQDM on the right arm of Chromosome III. We used RNA-mediated interference (RNAi) to test whether any of the coding genes in the QTL peak affected RQDM and found that *wah-1* was the sole gene to significantly affect RQDM. *wah-1* encodes the *C. elegans* orthologue of AIF1, a mitochondrial flavoprotein that affects the electron transport chain^26–29^, consistent with a potential role regulating RQDM. Finally, we showed that variation in WAH-1 activity is likely due to major differences in transcript structure between isolates and that there is similar complexity to *wah-1* transcripts in parasitic nematodes. We suggest that reduced expression of full-length WAH-1 protein results in more robust RQDM in both *C. elegans* and parasitic nematodes, and this identifies *wah-1* as a new regulator of anaerobic metabolism in these major human pathogens.

## Methods

### Worm strains and maintenance

All worm stocks were grown and maintained at 20℃ and were fed the *E. coli* strain OP50 on NGM agar plates as described elsewhere^30^. In addition to the traditional laboratory strain N2, wild *C. elegans* isolates used for genome-wide association have been described previously^22–24^, and were acquired from the *Caenorhabditis elegans* Natural Diversity Resource (CeNDR), including a set of 12 divergent strains (CB4856, CX11314, DL238, ED3017, EG4725, JT11398, JU258, JU775, LKC34, MY16, MY23, N2). The introgession lines (ILs) used for validating the QTL were graciously provided by Dr. Jan Kammenga’s group^31^. The CRISPR-engineered AF27 (*wah-1* E4 CAC indel in CB4856 background) strain was generated by SUNY Biotech.

### High-throughput Imaging Assays

All image-based assays were conducted on L1 animals as described previously^20, 21^. Briefly, L1 worms were collected by washing mixed-stage plates with M9 buffer and isolated using a 96 well 11 µM Multiscreen Nylon Mesh filter plate (Millipore: S5EJ008M04) to a final concentration of ∼120 animals per well. The worms were then incubated in varying concentrations of potassium cyanide (KCN; Sigma 60178-25G) for 15 hr, after which the KCN was diluted out 6-fold with M9 buffer. Immediately after dilution, worm movement was monitored every 10 min for 3 hr and quantified using an image-based quantification system as previously described^21^. All datapoints were normalized by the fractional mobility scores of the DMSO control wells per strain per time point.

### Preparation of the drugs for assays

KCN solutions were prepared fresh prior to each experiment by dissolving in phosphate buffered saline (PBS) and then diluting to a 5mM stock solution in M9 buffer. 2X working concentrations were then prepared with M9 and the KCN stock solution for a final concentration of 150 µM, 200 µM, 250 µM, and 300 µM. All of the wild isolates used for genome-wide association mapping were screened at 300 µM. All strains for all experiments contained 0.8% v/v DMSO control wells to account for strain-specific variation in movement prior to the administration of the drug.

Assays were assembled in flat-bottomed polystyrene 96-well plates (Corning 3997), with equal parts worms in M9 buffer and 2X KCN solution to a final volume of 40 µL per well.

### KCN dose-response assays

Dose-response experiments were performed in two divergent strains, N2 and CB4856, in technical triplicates prior to performing GWA experiments (Supp. Fig. 1A). Animals were assayed using the HTP image-based assay and analyzed as described above.

The KCN concentration (300 µM) chosen for GWA experiments was chosen based on an observable effect on worm movement phenotypes in the presence of KCN.

### GWAS using CeNDR and NemaScan

Genome-wide association mapping was performed on the *Caenorhabditis elegans* Natural Diversity Resource (CeNDR)’s web-based GWAS anaylsis pipeline, NemaScan^25^. All data used in the analysis was taken from the 20220216 CeNDR release^22^. ECA246 and ECA251 were removed from the mapping analysis due to the presence of duplicates from the same isotype group (CB4858, CB4853). The phenotypic trait input used for mapping was the fractional mobility score (FMS) at the 3- hour timepoint following recovery from KCN.

### RNAi assays

RNAi experiments were done by feeding as previously described^32, 33^. RNAi-treated and GFP RNAi control strains were filtered after 4 days to purify for L1 worms. The worms were washed with M9 buffer twice before being diluted to ∼120 worms per well in a 96- well plate and assayed using the HTP image-based assay described above.

### RNA-seq analysis of *wah-1* isoform expression in wild *C. elegans* isolates

Expression data on wild *C. elegans* isolates were reanalyzed from *Zhang et al, 2022*. Raw FASTQ reads were downloaded from the SRA under accession code PRJNA669810^34^. Three replicates for each isolate of interest were downloaded and aligned using HISAT2 (2.1.0)^35, 36^. SAM files were then sorted by coordinate order, converted to BAM format and the *wah-1* region was extracted using Picard (2.27.5). Total coverage and per-base depth for *wah-1* transcripts were calculated using SAMtools (1.16.1)^37^ and normalized to the mean depth of chromosome III. Coverage plots were generated using the mean of three replicates at each position. Manual review of sequencing reads was done using Integrated Genome Browser to determine the number of reads that splice through the Exon3-Exon4 junction.

### RNA-seq analysis of *wah-1* and *coq-2* isoform expression in clade V nematodes

We analyzed existing RNA-seq data for evidence of alternative isoforms of *wah-1* orthologues in parasitic nematodes. Publicly available RNA-seq data (ERP002173, SRP157940, ERP023010, SRP035476)^38–41^ were downloaded from the European Nucleotide Archive (ENA), with the accession numbers for the studies used listed in Supp. Table 3. Reads were first aligned to the parasite genome using HISAT2 using genome annotations taken from WormBase Parasite (WBPS17)^42, 43^. Reads were mapped to the PRJEB506 genome assembly for *H. contortus*^38, 44^, PRJNA231479 for *A. ceylanicum*^41^, PRJEB15396 for *H. polygyrus*^45^, and PRJEB511 for *N. brasiliensis*^45^.

Read coverage at each position of the *wah-1* orthologue gene locus and the averages across each chromosome/scaffold were then calculated using SAMtools on the resulting BAM files^37^. The number of reads at each position was then normalised to the average coverage across the entire chromosome/scaffold. For samples with more than one replicate, the mean normalised values were plotted. StringTie^46^ was then used for isoform quantification of *H. contortus wah-1* (HCON_00066460) and *coq-2* orthologues (HCON_00082210), using as input the BAM files from the HISAT2 alignments and a modified GTF transcript annotation file from WBPS17. We used the existing HCON_00066460 transcript annotations but manually added to the GTF file an additional isoform for HCON_00082210 corresponding to the COQ2e isoform that we previously predicted based on the available short-read sequencing data^47^.

## Results

### RQ-dependent metabolism varies in *C. elegans* natural isolates

Parasitic helminths—including roundworm, hookworm, and whipworm—infect over a billion humans worldwide and are able to spend extended periods in the highly anaerobic environment of the human gut^5–9^. To survive in this low oxygen condition, they use an unusual form of anaerobic respiration that relies on the alternative electron carrier rhodoquinone (RQ)^14–16^. RQ is highly related to ubiquinone (UQ) and is specifically utilized by these parasitic worms. Critically, RQ is not present in human hosts and understanding how RQ synthesis and use is regulated provides a promising avenue to identify new ways to target these parasites with drugs. It is challenging, however, to study parasites directly, but our group has previously shown that *C. elegans* is also capable of making RQ^19, 20^ and requires RQ-dependent metabolism (RQDM) to survive extended periods in potassium cyanide (KCN)^20^. This allowed us to establish a simple assay to measure RQDM in *C. elegans*. In brief, L1 larvae are placed in KCN for 18 hrs — the worms stop moving after ∼90 mins and remain immobile thereafter. This prolonged incubation with KCN triggers the switch from aerobic respiration to a metabolic state similar to that of parasitic helminths within their hosts, and their use of RQ under these conditions was hallmarked by the increased generation of succinate by complex II^48, 49^. The KCN is then diluted out with M9 buffer and the larvae then recover movement as measured using an image-based movement assay. However, worms carrying mutations that block RQ synthesis or treated with drugs that interfere with RQDM cannot survive the 18 hrs of KCN exposure^20^. The extent of recovery as read out by the movement phenotype thus gives us a quantitative readout of the ability of the worms to carry out RQDM.

We first tested a panel of 12 genetically diverse isolates in the published RQDM assay to determine whether there was variation in RQDM. The panel was chosen to cover a large amount of the known genetic variation in the *C. elegans* species. We found significant variation in the ability of the 12 isolates to recover from KCN-induced paralysis: CB4856, as well as some other highly divergent strains, exhibited more rapid and robust recovery compared to N2 (Fig. 1B, C). To test whether this difference in recovery appeared to be the result of variation on RQDM or whether the differences might be explained trivially by differing sensitivity to KCN itself, we tested whether N2 and CB4856 varied in the kinetics of initial paralysis following KCN exposure. We found no significant variation in the initial response to KCN between the two strains (Fig. 1A) with 300 μM KCN, suggesting that the variation seen in the ability to survive 18 hrs of KCN exposure was likely due to variation in their ability to carry out RQDM and not due to differences in acute KCN sensitivity.

**Figure 1:**
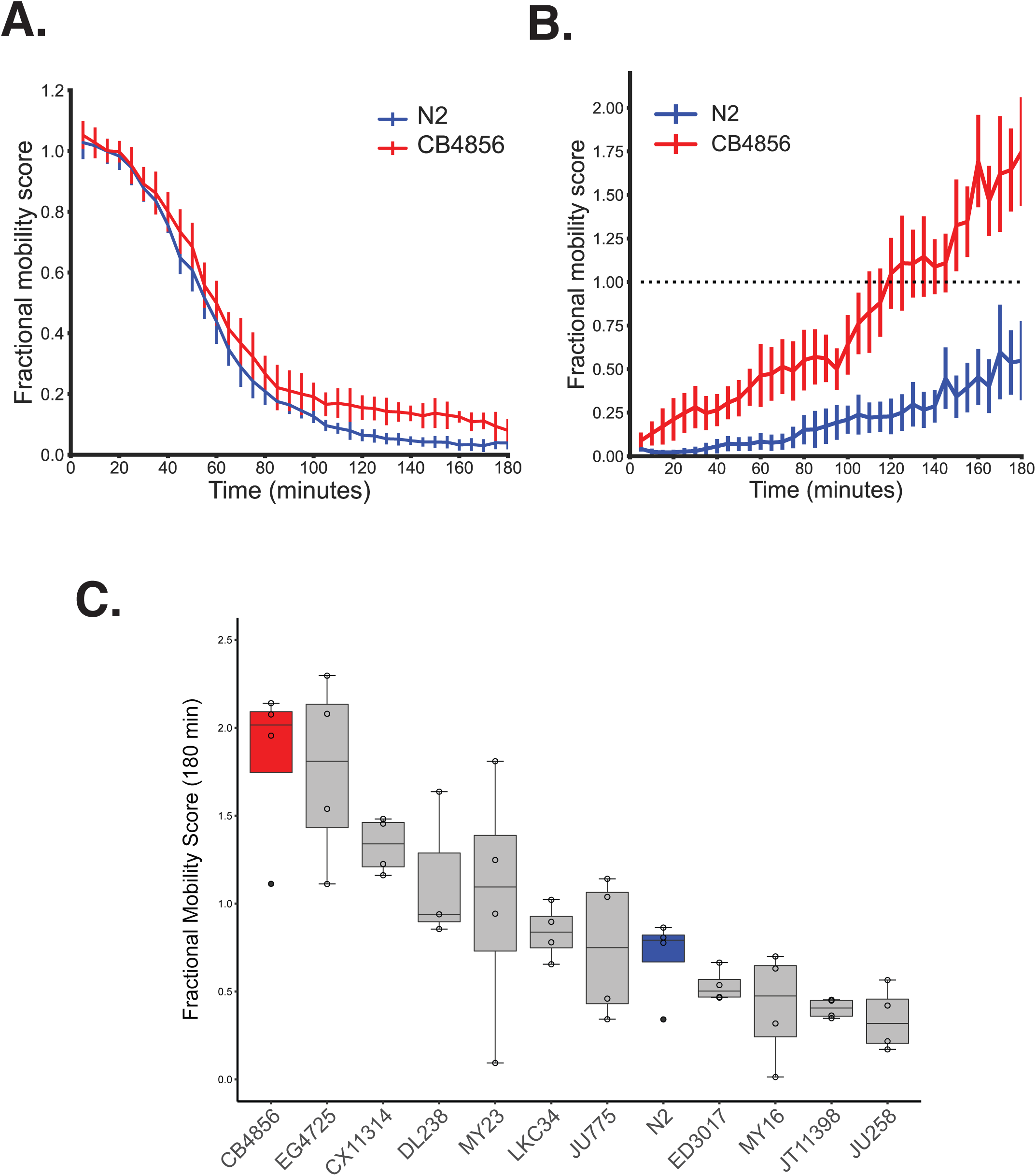
Natural isolates vary in their ability to survive exposure to potassium cyanide. A. Immediate effect of KCN treatment on worm movement does not vary between isolates: N2 and CB4856 L1 larvae were exposed to 300 µM KCN for 3 hrs and movement quantified using an image-based assay. Movement is expressed as a Fractional Mobility Score and is normalised to values for a no-drug (DMSO) negative control. B. Ability to survive long term KCN exposure varies between N2 and CB4856: N2 and CB4856 L1 larvae were exposed to 300 µM KCN for 18 hrs, and recovery from paralysis was measured following a 6-fold dilution with M9 for 3 hours. C. Ability to survive long term KCN exposure varies between isolates in a Diversity Panel: L1 larvae from 12 divergent isolates (CB4856, CX11314, DL238, ED3017, EG4725, JU258, JU775, JT11398, LKC34, MY16, MY23, N2) were treated with 300 µM KCN for 18 hrs, then the KCN was removed as before and movement quantified. Data are means of at least 3 independent replicates.

### Genome-wide association mapping identifies a single major QTL affecting RQDM

To map the QTLs that underlie the variation in RQDM in *C. elegans,* we tested 48 diverse isolates from the CeNDR collection (Fig. 2A) and performed a GWAS using the fractional mobility score (FMS) at 3 hrs post-recovery as our quantitative trait. This identified a single QTL on the right arm of chromosome III that passed the Bonferroni-corrected cutoff (Fig. 2B,C; Supp. Fig. 2A), whose peak at III:12340430 explains ∼40% of the phenotypic variance observed.

**Figure 2:**
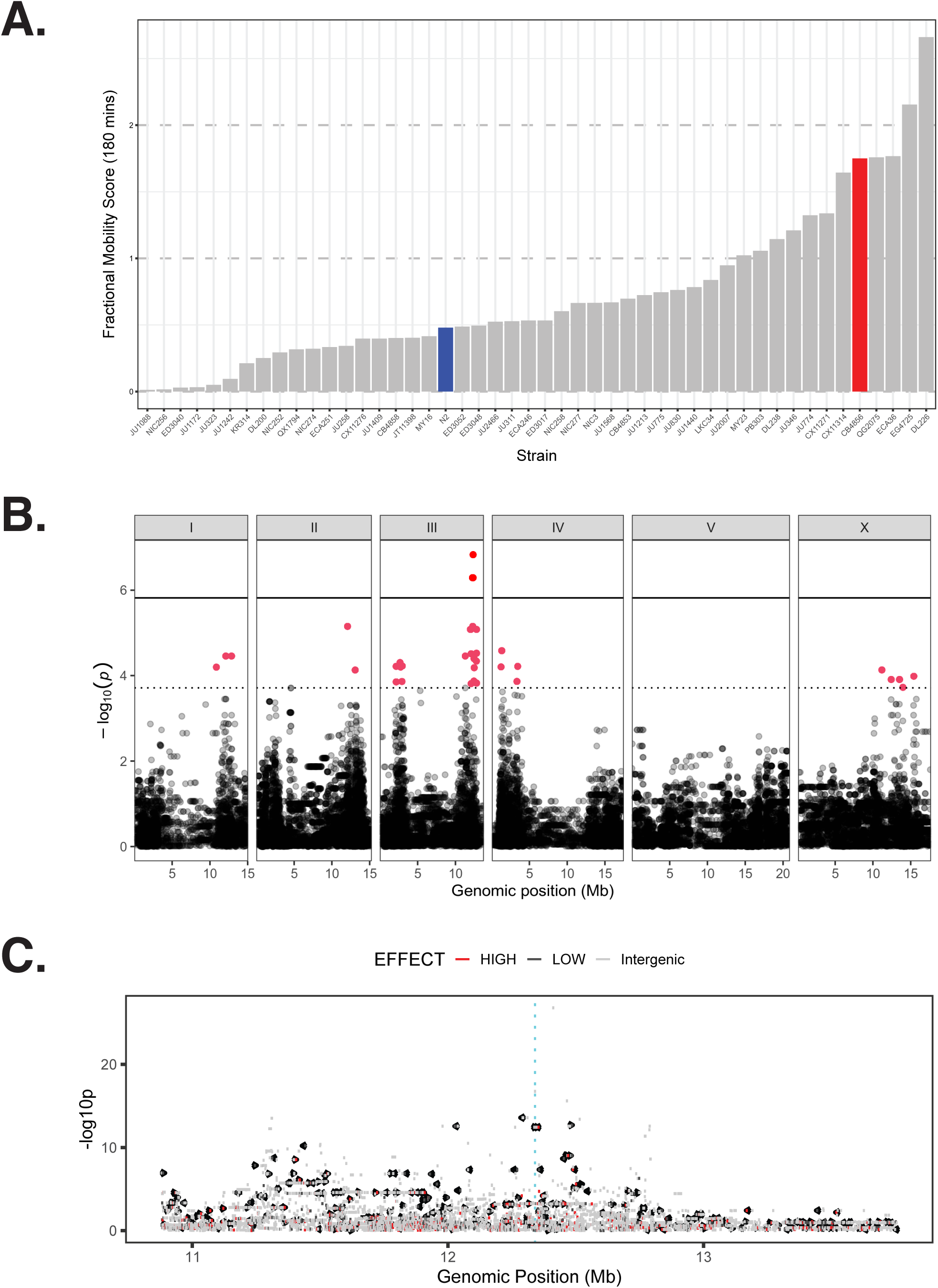
Variation in recovery from long term exposure to KCN is partly due to a QTL on Chromosome III. **A.** Barplot of the Fractional Mobility Scores (FMS) of L1 larvae from 48 wild *C. elegans* isolates were treated with 300 µM KCN for 18hrs before being diluted out with M9 and movement quantified after 3 hrs. Data are means of at least 3 independent replicates. **B. GWAS identifies a single significant QTL on the right arm of Chromosome III.** Manhattan plot of GWA mapping of the recovery from KCN treatment using NemaScan. FMS scores at 3 hrs post-KCN dilution was used as the quantitative trait. A single QTL on the right arm of chromosome III was found above the Bonferroni-corrected significance threshold (peak: III: 12,340,430). Each dot on the x-axis represents a SNV marker that is present in the assayed population at allele frequencies of at least 5%. The genomic position in Mb, separated by chromosome, is plotted on the x-axis and the *-log_10_(p)* for each SNV is plotted on the y-axis. Markers that pass either the Bonferroni-corrected threshold (solid line) or the genome-wide eigen-decomposition significance (ED) threshold (dashed line) are highlighted in red. **C. Fine mapping of the chromosome III region of interest**. SNVs located near the III:12340430 QTL were evaluated for their association with the phenotype. SNV markers coloured by predicted effects as defined using SnpEff.

The standard lab strain N2 and CB4856 show amongst the greatest differences in the RQDM assay. This was fortuitous since there are many tools previously established for fine mapping QTLs between these two strains. First, we wanted to test whether the variation in the region of chromosome III that was identified in our GWAS could explain the variation observed in the RQDM assay between N2 and CB4856. To do this, we screened a set of near-isogenic lines (NILs) from Doroszuk *et al,* 2009^31^. Each line within the collection contains a small interval of the CB4856 genome introgressed into the N2 genomic background, allowing us to assay small individual regions of the CB4856 genome in an otherwise N2 background. NIL strains were chosen to cover the entire QTL confidence interval, as well as the immediate left and right of the regions of interest (Fig. 3). We tested each NIL line in our RQDM assay and found that NILs containing sections of the CB4856 genome that covered the QTL peak on chromosome III had “CB4856-like” recovery. These results confirm that the locus identified on chromosome III contributes to the differences in RQDM between these strains and refined the interval containing the causal variant.

**Figure 3:**
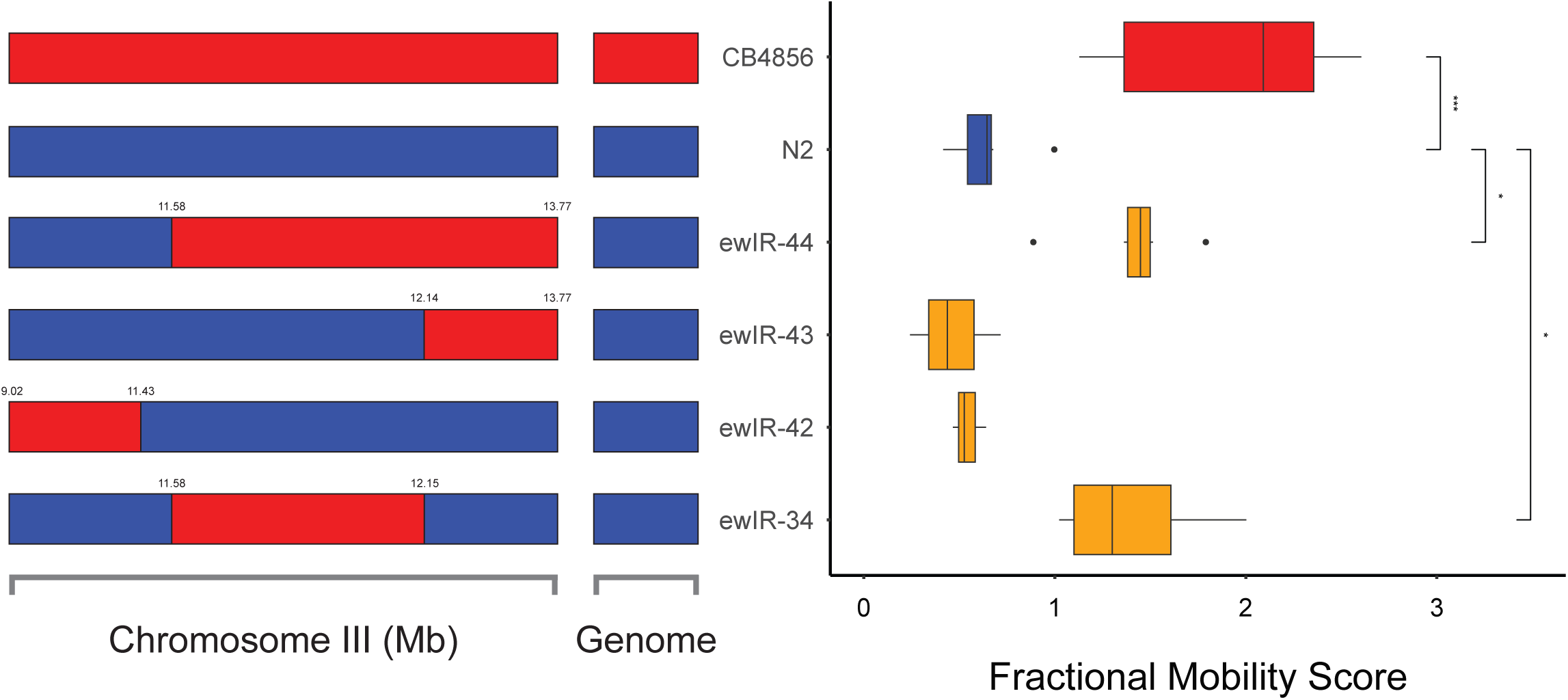
Nearly Isogenic Lines (NILs) map the QTL affecting KCN survival to a small region of Chromosome III. (Left) Strain genotypes are shown as colored rectangles (N2: blue; CB4856: red) in detail for chromosome III and for the rest of the chromosomes (Middle; “Genome”). (Right) FMS scores at 3 hrs (180 min) plotted as Tukey box plots against control (N2: blue & CB4856: red) and NIL strains (orange). Statistical significance of each NIL compared to N2 is calculated by Tukey’s HSD and is shown above each strain (N2-CB4856 = 0.0003, N2-ewIR-44 = 0.0156, N2-ewIR-34 = 0.396).

### RNAi knockdown screen of genes within the QTL interval identifies *wah-1* as affecting recovery from KCN

We identified a QTL on chromosome III that affected variation in RQDM in *C. elegans* isolates and used NILs to refine the region to 11.58 to 12.15 Mb. This interval contains 132 genes, 82 of which contain protein coding variants. To help identify the causal variant, we first chose to use RNA-mediated interference (RNAi) to identify which of the genes in the interval could affect RQDM. We aimed to test all 132 genes in the QTL interval — since interval mapping is not always precise, however, we also included genes in the regions flanking the immediate left and right boundaries of this interval, for a total of 196 candidate genes.

We carried out RNAi for each of the 196 genes using bacterial-mediated RNAi — this is robust in the N2 lab strain and results in reduced levels of the wild-type protein^32, 33^. For each gene, we placed L1 animals on a lawn of dsRNA-expressing bacteria and allowed them to grow to adulthood and lay the next generation. We purified L1s from the next generation and tested the L1 larvae in our RQDM assay — each gene was assayed three times in independent biological replicates. Their ability to recover after 18 hrs of KCN treatment was normalized to that of N2 worms treated with a GFP RNAi control, and z-scores were computed.

The N2 isolate shows relatively poor RQDM — other isolates like CB4856 show much more robust RQDM in our assay. We thus wanted to find if there were genes that altered the poor RQDM of N2 to the robust RQDM of CB4856 when targeted by RNAi — these would be candidates for the genes responsible for the variation in RQDM seen in the isolates. The sole significant effect was seen by targeting *wah-1* (Fig. 4A). Two independent RNAi clones were assayed for *wah-1* and both have the same result: N2 worms with reduced *wah-1* following RNAi have a greatly increased ability to recover after 18hrs of KCN treatment (Fig 4B), similar to the levels of recovery observed in CB4856. We conclude that variation of RQDM due to variation at the right arm of Chr III is likely due to variation in activity or expression of *wah-1* in our isolate panel. We then expanded our assays to test if the enhanced recovery following *wah-1* RNAi knockdown in N2 was seen in isolates that show similar RQDM phenotypes to N2. We find that isolates that show low “N2-like” recovery in the RQDM assay all show increased recovery following RNAi against *wah-1* (Fig. 5). We also note that RNAi against *wah-1* in isolates that show robust recovery like CB4856 have no significant effect on recovery, suggesting that the key difference between low “N2-like” recovery and robust “CB4856- like” recovery is indeed likely to be mainly due to reduced *wah-1* activity in CB4856-like isolates and this effect cannot be further enhanced by reducing expression further with RNAi.

**Figure 4:**
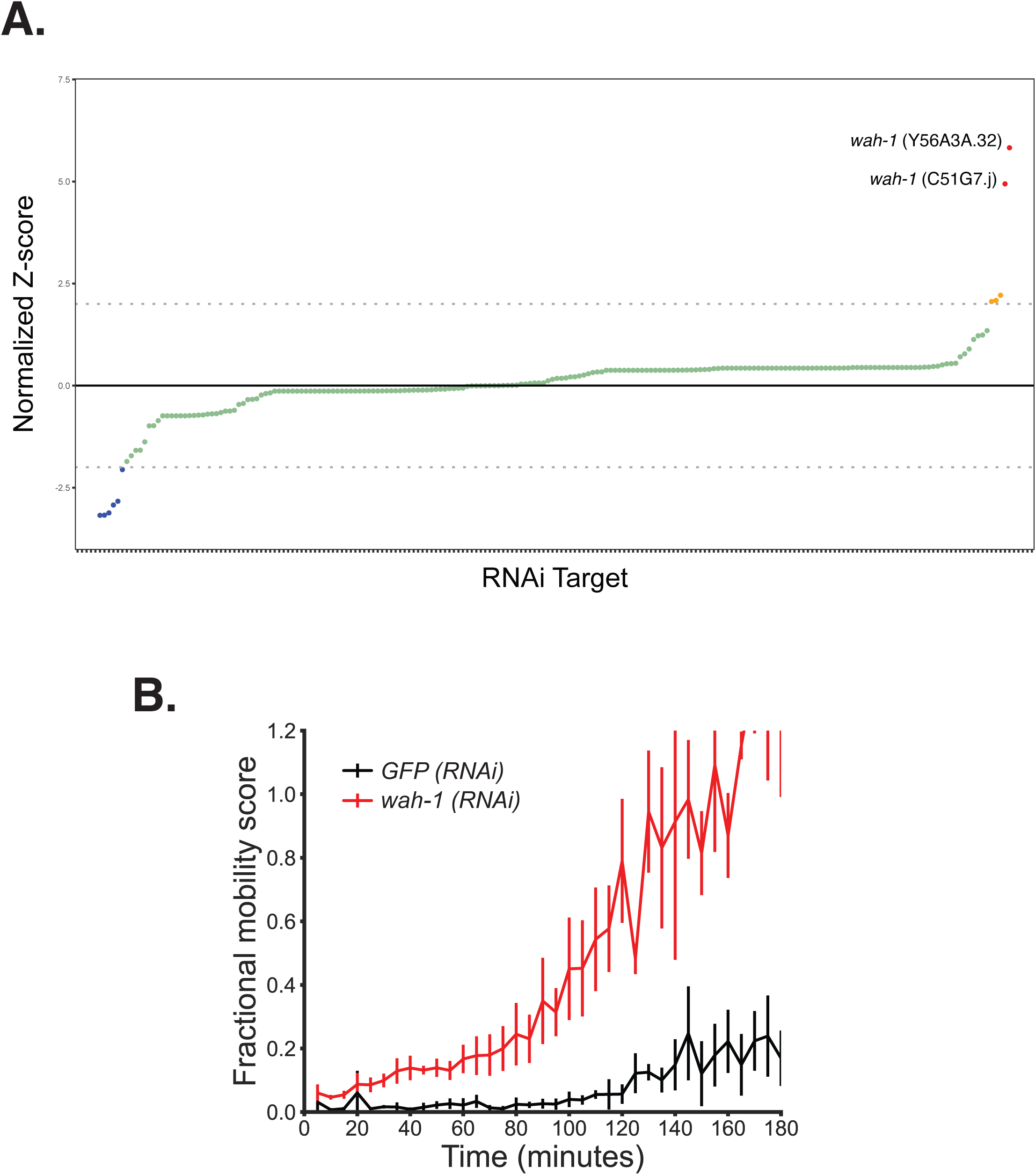
RNA-mediated interference (RNAi) identifies *wah-1* as the likely causal variant in the Chromosome III QTL interval. A. RNAi screening of genes on the right arm of chr III for impact on recovery following KCN exposure. N2 L1 worms were placed on lawns of dsRNA-expressing bacteria targeting 196 genes within the Chr III QTL interval and allowed to feed and develop for 96 hrs. At this point, the next generation L1 larvae were harvested and exposed to 200µM KCN for 18hrs. KCN was removed as described previously and worms were allowed to recover, and movement quantified after a further 3 hrs. The effect of RNAi of any specific gene is expressed as normalized Z-scores — this indicates how significantly the recovery differs to worms fed with dsRNA targeting GFP, a negative control. A significance cut-off of +2 and -2 (dashed lines) were used for targets whose knockdown results in better (red) or worse (blue) recovery compared to N2 controls, respectively. B. RNAi against *wah-1* results in enhanced recovery from KCN treatment. N2 L1 worms were placed on dsRNA-expressing bacteria targeting either *wah-1* (red) or GFP (black) as a negative control for 96 hrs. The next generation L1s were harvested and incubated with KCN for 18 hours before recovery. Time course of the recovery over 3 hours is shown. Data show means of three independent replicates.

**Figure 5:**
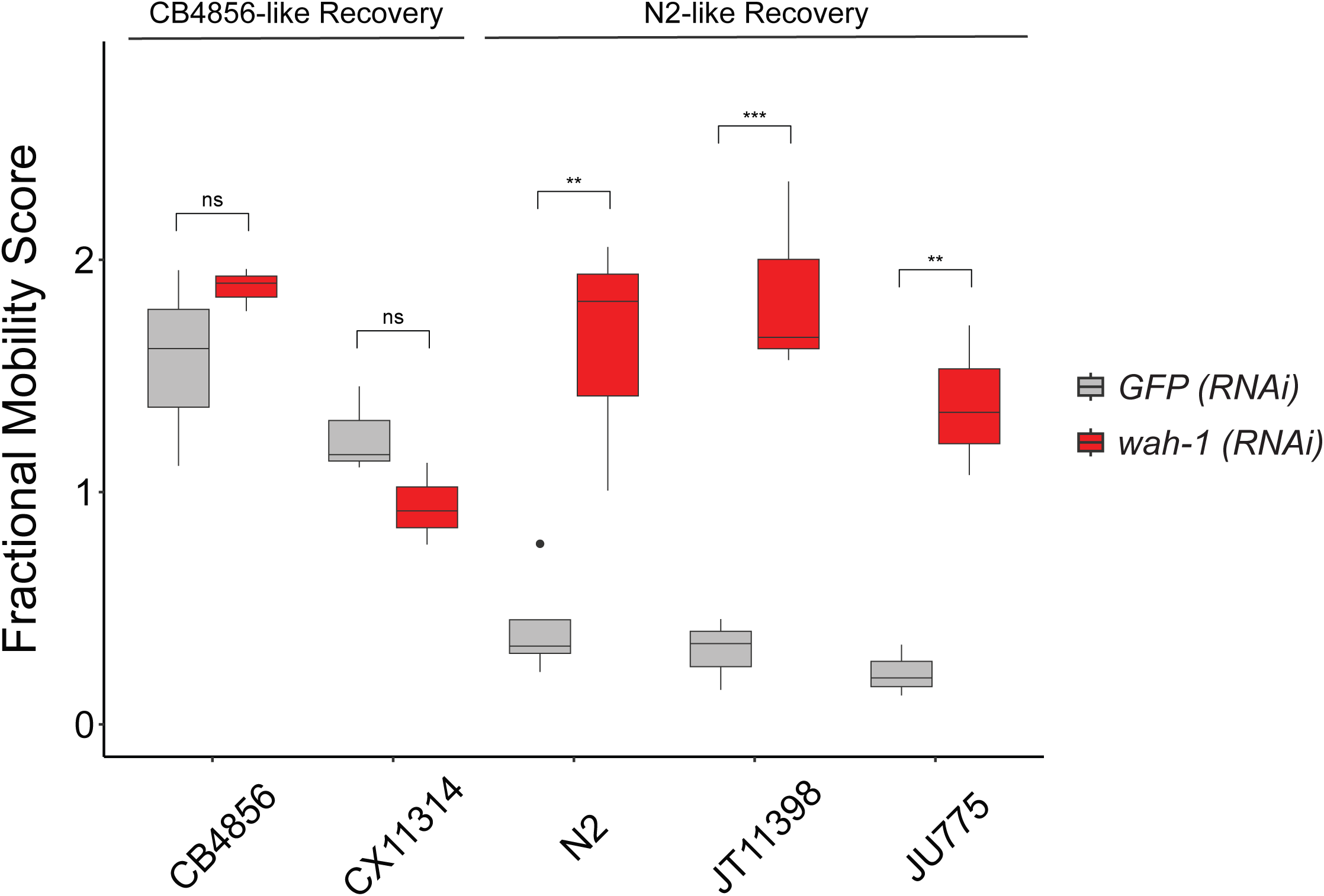
RNAi against *wah-1* results in increased recovery from KCN exposure in isolates that show weak recovery. Some isolates show robust recovery from 18 hrs of KCN like CB4856 and CX11314; others show weaker recovery like N2, JT11398 and JU775. L1 worms from each isolate were placed on lawns of dsRNA-expressing bacteria targeting either *wah-1* or GFP as a negative control for 96 hrs. At this point, the next generation L1 larvae were harvested and exposed to 200 µM KCN for 18hrs. The KCN was then removed, worms were allowed to recover, and movement quantified after a further 3 hrs. All pair-wise strain comparisons for N2-like recovering strains were significant (Tukey HSD p-values between GFP and *wah-1* RNAi treatments for each strain: CB4856 = 0.946, CX11314 = 0.961, N2 = 0.00117, JT11398 = 0.00012, JU775 = 0.004)

### Variation in *wah-1* activity is likely not the result of variation in WAH-1 coding sequence

WAH-1/AIF is a FAD-containing NADH-dependent oxidoreductase and is known to be a regulator of mitochondrial oxidative phosphorylation^26, 27^. It has previously been shown to be necessary for proper mitochondrial morphology and the biogenesis of various respiratory complexes, including Complex I, III and IV of the electron transport chain^29, 50^. Previous studies have also shown that decreased expression of *wah-1* lowers endogenous ROS levels, lowers complex I protein levels, and induces a nuclear-encoded mitochondrial stress response^51^. These functions are all consistent with a model that variation in activity of *wah-1* between isolates could impact the ability to do RQDM. To understand this in more detail, we looked at the coding variants in *wah-1* in the panel of isolates we had screened.

The CeNDR database contains both annotated coding variants for all isolates screened as well as the raw sequences files supporting those variant calls. We initially found that *wah-1* was annotated to show a very high degree of variation within its coding regions across isolates (CeNDR release 20170531), with multiple insertions and missense mutations across the whole panel. The extremely high number of coding variants made us suspicious of poorly aligned reads. Careful manual analysis of the BAM files for N2 and CB4856 revealed three synonymous mutations located on exons 6 and 8, as well as an unannotated insertion of a trinucleotide CAC resulting in the insertion of an additional threonine near the beginning of exon 4. Isolates containing this CAC insertion tended to recover better than N2 (Fig. 6A). To test if this mutation was indeed causal for the hyper-recovering phenotype, we generated a CB4856 strain in which the CAC insertion found in the hyper-recovering isolates was deleted to see if it recapitulates the N2-like recovery response. However, this engineered strain showed no difference in our RQDM assay to CB4856 (Fig. 6B). We therefore conclude that the additional threonine that results from a CAC expansion in CB4856 is not the cause of the enhanced RQDM compared with N2 and that there is no consistent coding difference in the *wah-1* gene that can explain the variation in RQDM assay between isolates. We therefore examined if *wah-1* expression varied between isolates.

**Figure 6:**
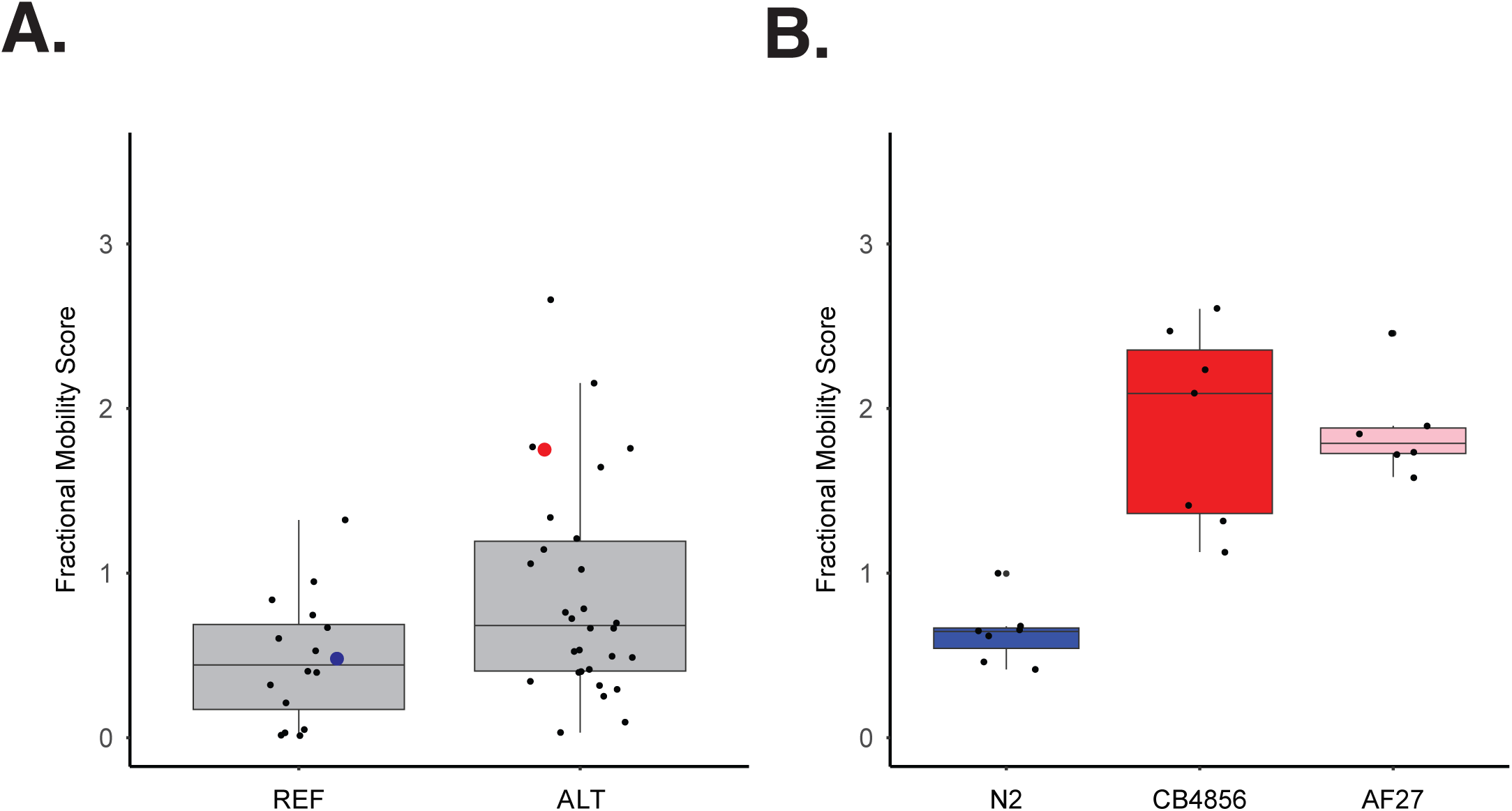
The deletion of the unannotated CAC insertion near the beginning of exon 4 on *wah-1* has no effect on CB4856 recovery. A. Genotype-phenotype Tukey box plots of the fractional mobility scores used for association mapping. Each dot represents the phenotype of an individual strain in the assayed population, as plotted on the y-axis. Strains are grouped by their phenotype at position III:11,994,425, wherein REF corresponds to the allele from the laboratory N2 strain and where ALT refers to the CAC insertion that occurs in CB4856. N2 highlighted in blue, CB4856 highlighted in red. B. Deletion of the CAC insertion in the CB4856 background has no effect on CB4856 recovery. Tukey box plots of N2, CB4856, and a mutant strain containing the deletion of the CAC insertion was generated in the CB4856 background (AF27; pink). Strains were treated with 300 µM KCN for 18hrs and allowed to recovery for 3 hrs. Statistical analysis with Tukey’s HSD showed no significant difference between AF27 and CB4856.

### *wah-1* transcript structures vary between natural isolates and this correlates with variation in RQDM phenotypes

Since we could not identify coding differences that correlate with the observed variation in RQDM, we examined variation in *wah-1* expression among the assayed isolates. We used previously published RNA-seq data to compare three strains that showed “N2-like” KCN response (N2, JT11398 and JU775) with strains that had the less common “CB4856-like” response (CB4856, CX11314 and QG2075) to determine if there were consistent expression differences in *wah-1* in these two groups of strains.

Superficially, expression of *wah-1* appears the same in all 6 isolates — the overall RPKM values from RNA-seq are very similar and thus there seems to be no variation in gene expression. Closer inspection of the transcripts, however, showed that they appeared mis-annotated on Wormbase (WS287) and this turned out to be critical. *wah- 1* has 8 exons and in N2 there is high transcription over exons 1-3 and lower transcription over exons 4-8 (Fig. 7A; blue track). This appears to be because there are two alternative 3’ ends to *wah-1* transcripts in N2. One transcript is the correctly annotated *wah-1, isoform a* transcript which runs from exon 1 (E1) to E8 and encodes full-length WAH-1a. The other *wah-1* transcript is short, including only exons 1-3 then extending into intron 3 and ending near a consensus AAUAAA poly(A) signal in intron 3 — this transcript is not correctly annotated and we propose to name this *wah-1, isoform d*. Consistent with this, nanopore sequencing of N2 shows a transcript that runs from exon 1 through to this poly(A) site in intron 3 and 3′ end sequencing also shows a poly(A) cluster in this intron^52^.

**Figure 7:**
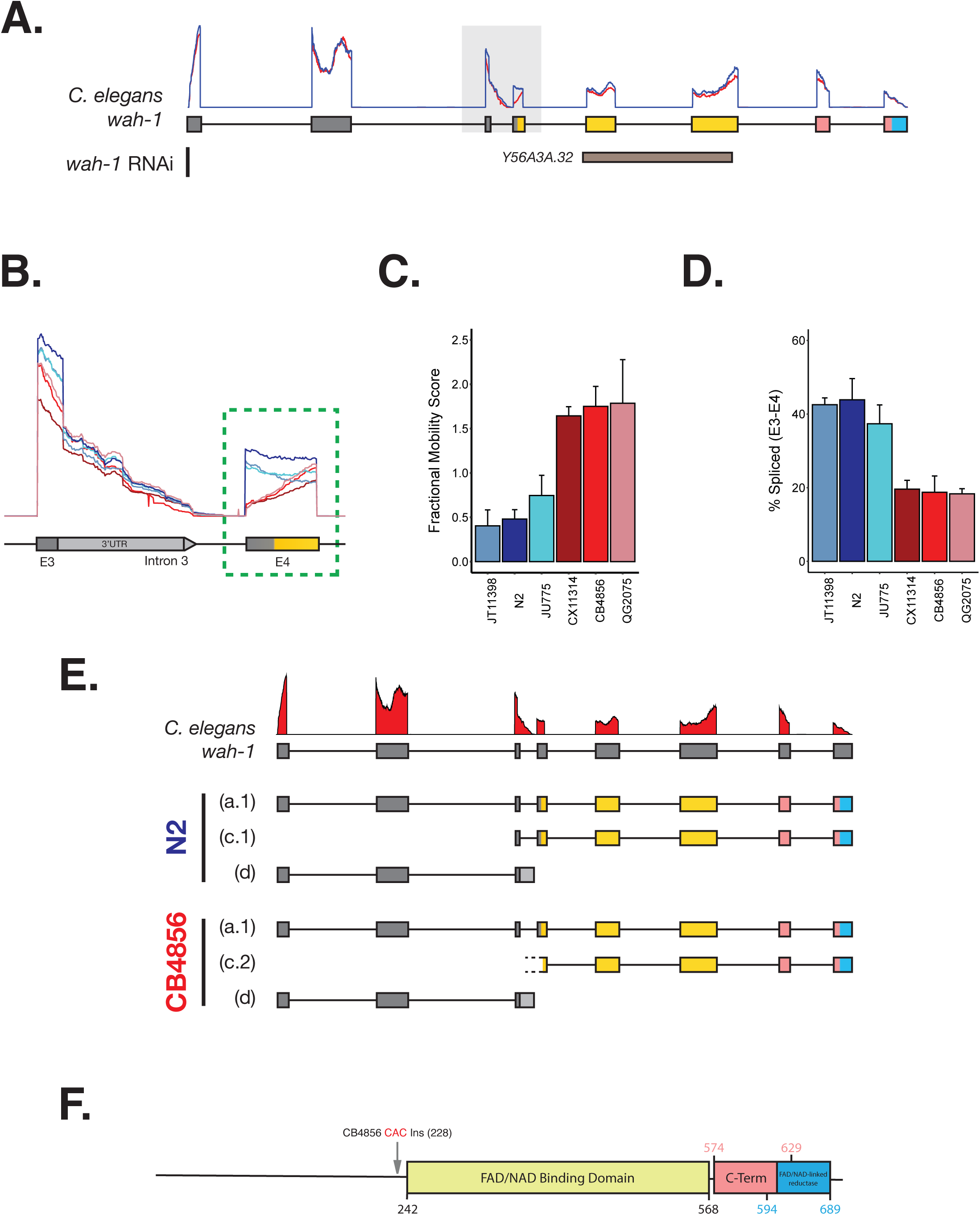
*wah-1* transcript structures vary between isolates at the E3-I3-E4 splice junction and correlates with variation in recovery from KCN. A. Full-length *wah-1* transcript structure. Top: coverage plot of N2 (blue) and CB4856 (red) RNA-seq data. Coverage depth normalized to the mean depth of chr III. Bottom: Location of the sequence targeted by the *wah-1* RNAi clone (Y56A3A.32). B. Zoomed-in coverage plots of the Exon3-Intron3-Exon4 junction. Coverage plots for *wah-1* from three isolates with N2-like recovery and three isolates with CB4856-like recovery. RNA-seq data were extracted, and coverage depth for each strain was normalized to the mean depth of chr III. Data represents the average depth at each position of three independent replicates. C. Fractional mobility scores of the six isolates at 3 hours post-recovery from KCN. Isolates with N2-like transcript structure highlighted in shades of blue; isolates with CB4856-like transcript structure highlighted in shades of red. Data is mean of at least four independent replicates. D. Isoform estimations derived from normalized depth of coverage at the end of Exon 3 vs start of Intron 3. Marker used for E3 - III:11994103; marker used for I3 – III:11994104. Percent spliced was calculated as the difference of coverage between E3 and I3 divided by the total coverage over the junction. Data is mean of three independent replicates. E. Proposed *wah-1* transcript structures in N2 and CB4856. Exon colours correspond to the predicted protein domains in (7F) below. F. Predicted protein domains in N2 WAH-1. Yellow = FAD/NADH binding domain; pink = C-terminal domain; blue = FAD/NAD-linked reductase domain. Location of the CAC insertion found in CB4856 highlighted in red.

Quantifying exon junction counts indicates that in N2, about half of the transcripts are the short, unannotated E1-3 *wah-1d* transcript that ends in intron 3, while the other half of the transcripts are full-length E1-E8 *wah-1a* that include splicing from exon 3 to exon 4. This is essentially the same picture in all 3 isolates that show “N2-like” RQDM (Fig. 7B,C). The picture is very different in the isolates with “CB4856-like” RQDM (CB4856, CX11314 and QG2075). These strains all show strong transcription of the short E1-E3 *wah-1d* transcript, similar to the "N2-like” isolates — however there is very little splicing from E3 to E4 (Supp. Table 1-2). There is thus a greatly reduced level of full length *wah- 1a* E1-E8 transcripts in the isolates with “CB4856-like” RQDM.

In “CB4856-like” isolates, the great majority of transcripts that enter E3 are not spliced to E4 but end in intron 3 at the alternative polyadenylation site, making the novel *wah- 1d* transcript. If most CB4856 *wah-1* transcripts end in intron 3, this suggests there should be a lower level of transcription of exons 4-8 in CB4856 compared with N2. Remarkably, however, exons 4-8 are transcribed at a similar level as in N2. This appears puzzling — where are these transcripts in E4-E8 initiating? In “CB4856-like” isolates, the E4-E8 transcription appears to be due to novel transcriptional initiation in CB4856 within exon 4 (Fig. 7C; isoform c.2). The read coverage ‘ramps up’ across E4 in all “CB4856-like” isolates — the standard signature of transcriptional initiation — and we note that there is an annotated peak of RNA polII occupancy in intron 3. We also note that there is an in frame ATG near the start of this novel transcript, suggesting that it is indeed a coding transcript that encodes the critical FAD and NAD(H)-binding domains of WAH-1 (Fig. 7F). Thus, while “N2-like” isolates appear to make ∼40:60 mix of E1-E3 short *wah-1d* and full length *wah-1a,* “CB4856-like” isolates make >60% E1-E3 short *wah-1d, ∼*20% full length *wah-1a,* and the remainder is this unique *wah-1* transcript initiating in E4 and extending to the normal 3’ end of *wah-1* after E8. These transcripts are summarized in Fig. 7E.

What are the consequences of the different transcripts made in different isolates on the WAH-1 proteins made in N2-like isolates and CB4856-like isolates? The first and most obvious is that there is a strong difference in the level of full-length *wah-1a* E1-E8 transcript and thus likely in the level of full-length WAH-1a protein. CB4856 makes ∼2x less full-length *wah-1a* transcript than N2 and recovers better from extended KCN exposure, which is consistent with the finding that reducing the level of WAH-1a in N2 using RNAi results in enhanced recovery. In addition to the difference in levels of full-length WAH-1a, there are also key differences in expression of the short forms of WAH-1. All isolates examined make similar levels of the E1-E3 transcript and are thus likely to make similar amounts of this N-terminal short WAH-1d. However, only the “CB4856- like” isolates make significant levels of the truncated E4-E8-encoded C-terminal short WAH-1 that includes the FAD and NAD(H)-binding domains. There is considerable complexity and variation in the transcript structures at the *wah-1* locus, and thus in the proteins made, and *wah-1* transcription varies strongly between isolates that show poor recovery from extended KCN treatment and isolates that have more robust recovery.

### Other Clade V nematodes show complex transcription of *wah-1*

We have found differences in *wah-1* transcription between isolates of *C. elegans* that show robust RQDM (“CB4856-like”) and those that have reduced RQDM (“N2-like”) — lower expression of full length *wah-1a* correlates with more robust RQDM. This suggests that regulation of *wah-1* expression may have biological importance in situations where rhodoquinone-dependent metabolism (RQDM) is critical. We thus asked whether changes in *wah-1* expression might impact RQDM in other nematodes where it is critical for their survival. Parasitic nematodes such as hookworms and whipworms require RQDM to survive in the highly hypoxic environment of the host gut^7, 8^. Since changes to *wah-1* expression affect how well *C. elegans* isolates can survive in conditions where they rely on RQDM, we thus wondered whether parasitic nematodes might regulate *wah-1* transcription in a similar manner.

We examined the transcriptomes of Clade V parasites *H. contortus*^38^, *H. polygyrus*^39^, *N. brasiliensis*^40^*, and A. ceylanicum*^41^, and found similar complexity to *wah-1* transcription as in *C. elegans*. Like N2, the Clade V worms seem to make two main transcripts — a short 5’ transcript that corresponds closely to the short E1-E3 *C. elegans wah-1d* transcript, and a full-length transcript similar to *wah-1a* (Fig. 8A). Interestingly, we see a consistent pattern in the shorter transcripts, wherein they extend into the intronic region just before the splice site (Fig. 8B), and like in *C. elegans*, the functional FAD/NAD(H)- binding domains is specifically encoded by the full-length transcript (Supp. Figure 3).

**Figure 8:**
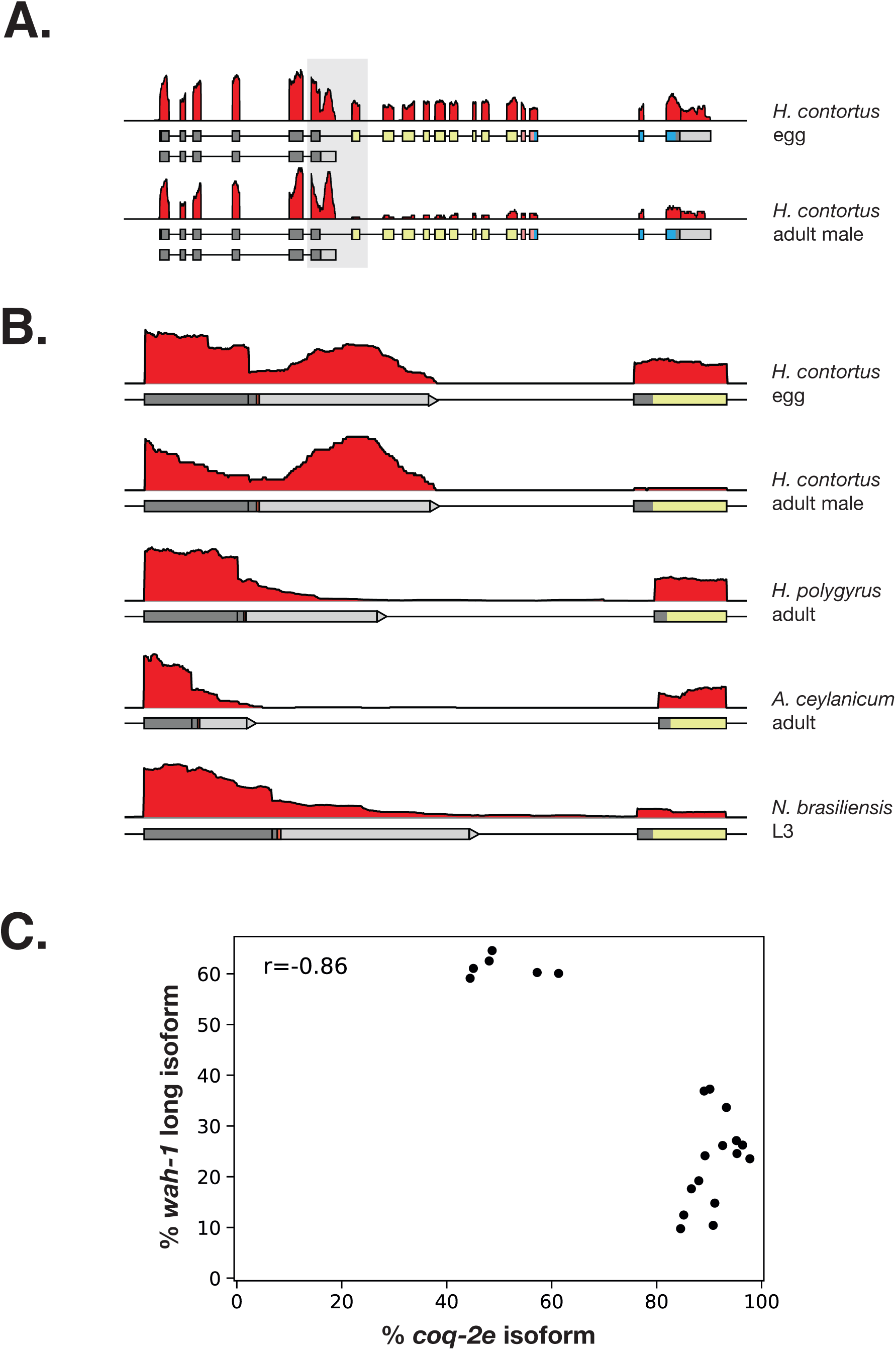
Parasitic nemetodes also show complex transcription of *wah-1*. A. Coverage plots for the *wah-1* orthologue in *H. contortus*. RNA-seq data from adult males and eggs were extracted from *Laing et al, 2013* and normalized to the average of the entire chromosome. Coverage for *wah-1* transcripts in both strains were then averaged across of three biological replicates. Exon colours correspond to the predicted protein domains as found in Supplementary Figure 3. **B. Zoomed-in coverage plots for the Exon3-Intron3-Exon4 junction across other clade V nematodes.** Data represents the average of three biological replicates for all except for *A. ceylanicum* (one biological replicate). **C. *wah-1* vs *coq-2e* scatterplot**. Increased inclusion of the RQ-specific exon in *coq-2* (HCON_00082210; unannotated isoform *’coq-2e’*) is correlated with decreased proportion of the long *wah-1* isoform (HCON_00066460-00001) relative to the short isoform (HCON_00066460-00002). Isoform-level expression of *wah-1* and *coq-2* at different stages of the *H. contortus* life cycle was calculated using StringTie on data derived from public RNA-Seq data (Laing et al, 2013) (Further details found in Methods and Supplementary Table 3).

As parasites switch from aerobic to anaerobic conditions, they alter their quinones from majority UQ (aerobic) to majority RQ (anaerobic). We previously showed that the critical switch from UQ to RQ synthesis is an alternative splicing event in the polyprenyltransferase COQ-2^47^. This splicing switch from COQ-2a (UQ synthesis) to COQ-2e (RQ synthesis) occurs as parasites move from aerobic to anaerobic conditions and high COQ-2e expression can thus be used as a proxy for use of RQDM. We therefore examined whether *wah-1* expression was related to levels of RQDM by comparing *wah-1* expression with *coq-2e* expression. We find that, across the various developmental stages in *H. contortus*, where there is rich transcript data across the full life cycle, there is an inverse relationship between the levels of the *coq-2e* isoform and full-length *wah-1* transcripts. In developmental stages with a high abundance (∼80- 100%) of the *coq-2e* isoform (i.e. relying on RQDM), there is less of the long *wah-1* isoform (Fig. 8C). However, in the stages with lower *coq-2e* (and thus less RQDM) there is a higher proportion of the long *wah-1* isoform (∼60%). Expression levels of full-length WAH-1 thus change in parasitic nematodes in a way that is consistent with a role in anaerobic RQ-dependent metabolism. During periods where parasites rely on RQDM, they express low levels of full-length WAH-1 and instead generate mainly shorter transcripts that encode truncated versions of WAH-1, suggesting that low levels of full-length WAH-1 are likely beneficial in conditions where nematodes rely on RQDM. This is similar to what we observe in natural isolates of *C. elegans* — isolates with lower levels of full-length *wah-1* due to alternative transcript expression survive better under conditions where they rely on RQDM for survival, and reducing *wah-1* expression results in better survival in RQDM-requiring conditions. We conclude that differences in expression of full-length WAH-1 impact the ability of both free-living and parasitic nematodes to survive in conditions where they require on RQDM.

## Discussion

Nematodes can survive in highly hypoxic conditions by using an unusual form of anaerobic metabolism that relies on the quinone electron carrier rhodoquinone (RQ)^7, 14–16^. We previously showed that survival of *C. elegans* in potassium cyanide (KCN) requires RQ-dependent metabolism (RQDM) and established a simple movement-based assay for RQDM^20^. Strains that can carry out RQDM survive 18hrs of treatment with KCN and recover movement following KCN withdrawal, while strains with reduced RQDM capacity show weaker recovery, and RQDM defective strains are all dead after prolonged incubation with KCN. The extent of recovery after 18hrs of KCN exposure is thus a simple readout for ability to carry out RQDM. Here, we examined a collection of natural isolates of *C. elegans* to see if there was natural variation in the ability to carry out RQDM using this assay.

We found that most isolates have similar responses to the lab strain N2, showing moderate recovery of movement after 18hrs of KCN exposure. However, a small number of isolates, including the divergent strain CB4856, show a much greater recovery in our RQDM assay, indicating an enhanced ability to do RQDM. We used GWAS to identify one of the major QTLs responsible for this variation in RQDM — this maps to the right arm of Chromosome III and explains ∼40% of the variance in the RQDM response. The validated QTL peak covered ∼130 genes and we used RNAi to test whether reducing expression of each gene in the interval could affect RQDM. We found a single gene in the QTL interval that affected RQDM, *wah-1*. This encodes a mitochondrial flavoprotein that is known to affect levels of several complexes of the electron transport chain (ETC), including Complex I^29, 50^. It is likely to act as an alternative quinone-coupled entry point for electrons into the ETC like Ndi1 in *S. cerevisiae*^53, 54^. The biochemical activity of WAH-1 is thus consistent with a possible effect on RQDM which is fundamentally a rewiring of the ETC.

We found no consistent *wah-1* coding change that appears responsible for the variation in RQDM across the natural isolates. Instead, we found a remarkable difference in transcript structures between isolates that show moderate recovery (“N2-like”) and strong recovery (“CB4856-like”) from 18hrs in KCN. N2-like strains make two main transcripts — one includes exons 1-8 in a spliced transcript that encodes a full-length WAH-1 protein and is the annotated *wah-1a* transcript. The other is a short transcript that includes exons 1-3, but instead of splicing from exon 3 to exon 4, it transcribes into intron 3, resulting in a short unannotated *wah-1* transcript that we propose to call *wah- 1d*. This would only encode an N-terminal portion of WAH-1 that lacks the FAD and NAD binding domains, and it is unclear what the function of this N-terminal portion of WAH-1 could be. We note however, that the RNAi clones that we used to target *wah-1* would not affect this *wah-1d* transcript (Fig. 7A) — the involvement of WAH-1 in RQDM that we observe must thus either be due to full length WAH-1a or the novel C-terminal form of WAH-1 seen in “CB4856-like” isolates. We have, thus far, no evidence for any involvement of WAH-1d in RQDM.

N2 and other “N2-like” strains make almost entirely *wah-1a* and *wah-1d* in a ∼40:60 ratio. CB4856 and other strains that also robustly recover from KCN, on the other hand, make similar levels of the short *wah-1d* transcript as N2 but make strikingly much less full-length *wah-1a*, with only ∼20% of the transcripts showing splicing from exon 3 to exon 4. The result is that CB4856 makes ∼2-fold less full length *wah-1a* compared to N2-like strains, and this is consistent across all “CB4856-like” isolates we examined. More remarkably, “CB4856-like” strains appear to make a new transcript that is absent from “N2-like” strains — they have a new transcript starting within exon 4 upstream of an in-frame start ATG codon, including E4 and all downstream exons. This novel CB4856 transcript is not found in N2 even by long-read sequencing^55^, and would encode a short form of WAH-1 that includes the key FAD and NAD binding domains. We note that this novel C-terminal WAH-1 corresponds closely to specific isoforms of mammalian AIF, the orthologue of WAH-1^56, 57^. AIF is thought to be cleaved in mitochondria in its N-terminal region, releasing a C-terminal region that contains the FAD and NAD binding regions, and this released form is very similar to the short C-terminal form of WAH-1 only seen in “CB4856-like” isolates. We currently have no model to explain the difference in transcript structure between “N2-like” and “CB4856- like” isolates. We see no consistent variation in exon 3, exon 4 or in intron 3 that might explain this and thus conclude that the cause is unlikely to be due to variation in a *cis-* acting element and is more likely due to some more complex *trans* effect.

The transcript differences we see between “N2-like” and “CB4856-like” isolates results in major differences in WAH-1 proteins. The critical difference is likely to be that CB4856-like isolate make ∼2-fold less full-length WAH-1a protein since they fail to splice efficiently from E3 to E4. The lower level of full length WAH-1a in CB4856-like isolates is consistent with our RNAi data: we found that reducing *wah-1* levels in N2 results in CB4856-like recovery. These data all suggest that lower *wah-1a* expression in CB4856-like strains due to variation in transcript structures results in enhanced ability to carry out RQDM and thus survive extended exposure to KCN.

To gain some insight into whether changes in *wah-1* transcription might impact RQDM- relevant biology, we turned to parasitic helminths. These major pathogens rely almost entirely on RQDM when they are in the host gut, which is highly anaerobic. We found that other clade V nematodes such as *H. contortus*, *H. polygyrus*, and *N. brasiliensis* all show similar alternative polyadenylation in *wah-1*. All parasites examined appear “N2- like” in the transcripts: they make a short *wah-1d*-like transcript and a full-length *wah-1a* transcript. Crucially, in *H. contortus*, the parasites appear to switch from making both *wah-1a* and *wah-1d* at stages when they make a mix of both the *coq-2a* and *coq- 2e* isoforms to making almost exclusively short *wah-1d* transcripts in conditions where they are largely making only the *coq-2e* isoform that is specific to RQ-producing animals (Fig. 8C). The correlation of increased inclusion of the RQ-specific exon in COQ-2 with lower expression of full-length WAH-1a thus suggests lower expression of WAH-1a might be important for RQDM use in parasites, and this is consistent with our finding that reduced WAH-1a levels—either as the result of natural variation or RNAi— results in more robust RQDM in *C. elegans*. We conclude that variation in *wah-1* transcript structures between isolates results in variation in RQDM in *C. elegans,* and that similar changes in *wah-1* transcript structures occur in parasites in a way that is consistent with a modulating role for WAH-1 in RQDM.

## Acknowledgements

This research was funded by grants PJT-153036 and PJT-148637 to Dr. Andrew Fraser from the Canadian Institute of Health Research (CIHR). We would like to thank Dr. Erik Andersen for worm strains and for his insightful guidance with regards to the CeNDR platform; Dr. Jan Kammenga for the NIL strains; as well as Dr. Asher Cutter, Dr. Will Ryu, Dr. Peter Roy, and the rest of the Fraser lab for their helpful discussions throughout.

---

**Supplementary Figure 1.**
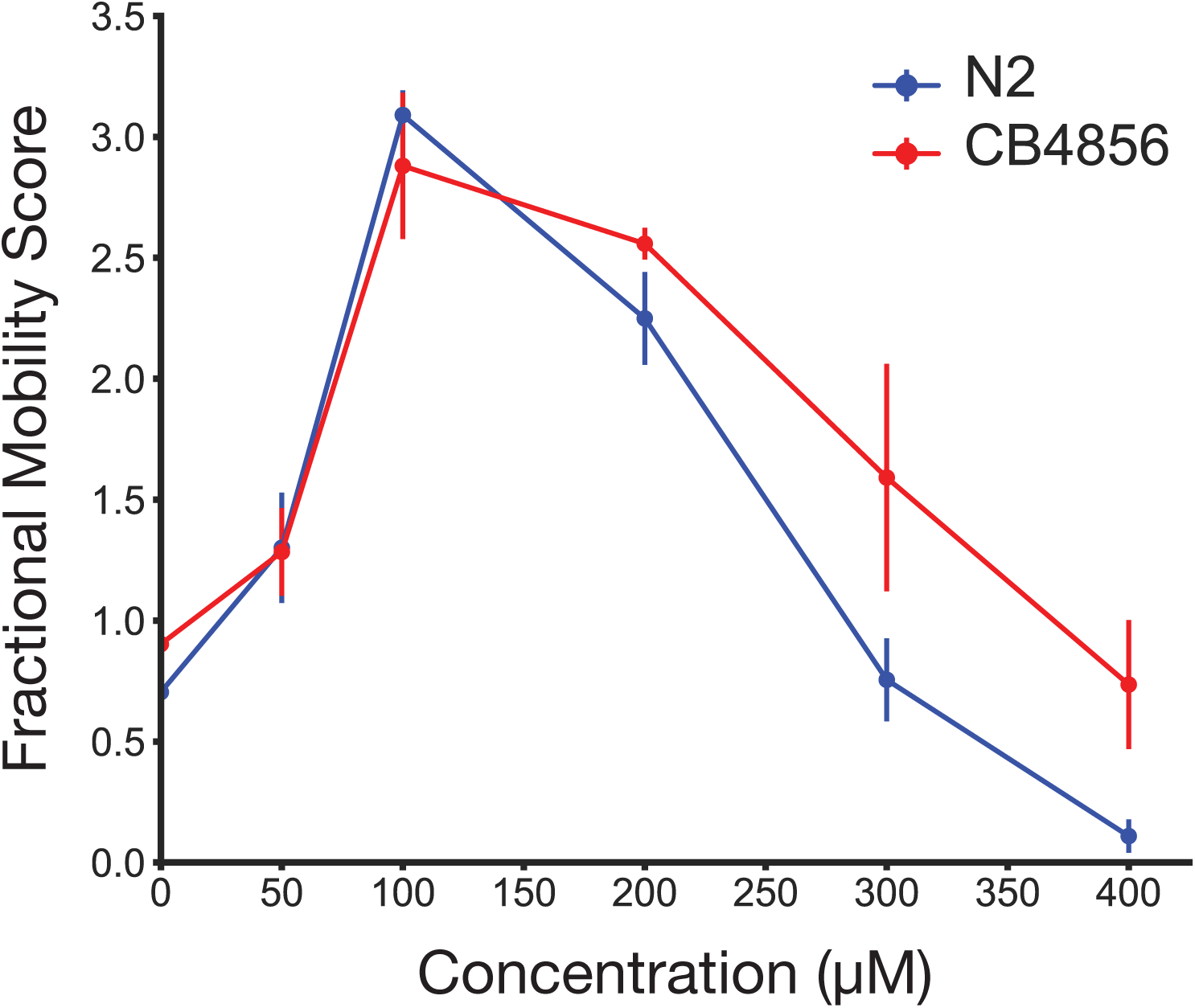
Dose-response analysis of N2 (blue) and CB4856 (red) against 50, 100, 200, 300 and 400 μM KCN. L1 larvae from each strain was treated with KCN for 18 hrs, and movement was captured at 3 hrs post-M9 dilution of KCN. Data represents the mean of three biological replicates.

**Supplementary Figure 2.**
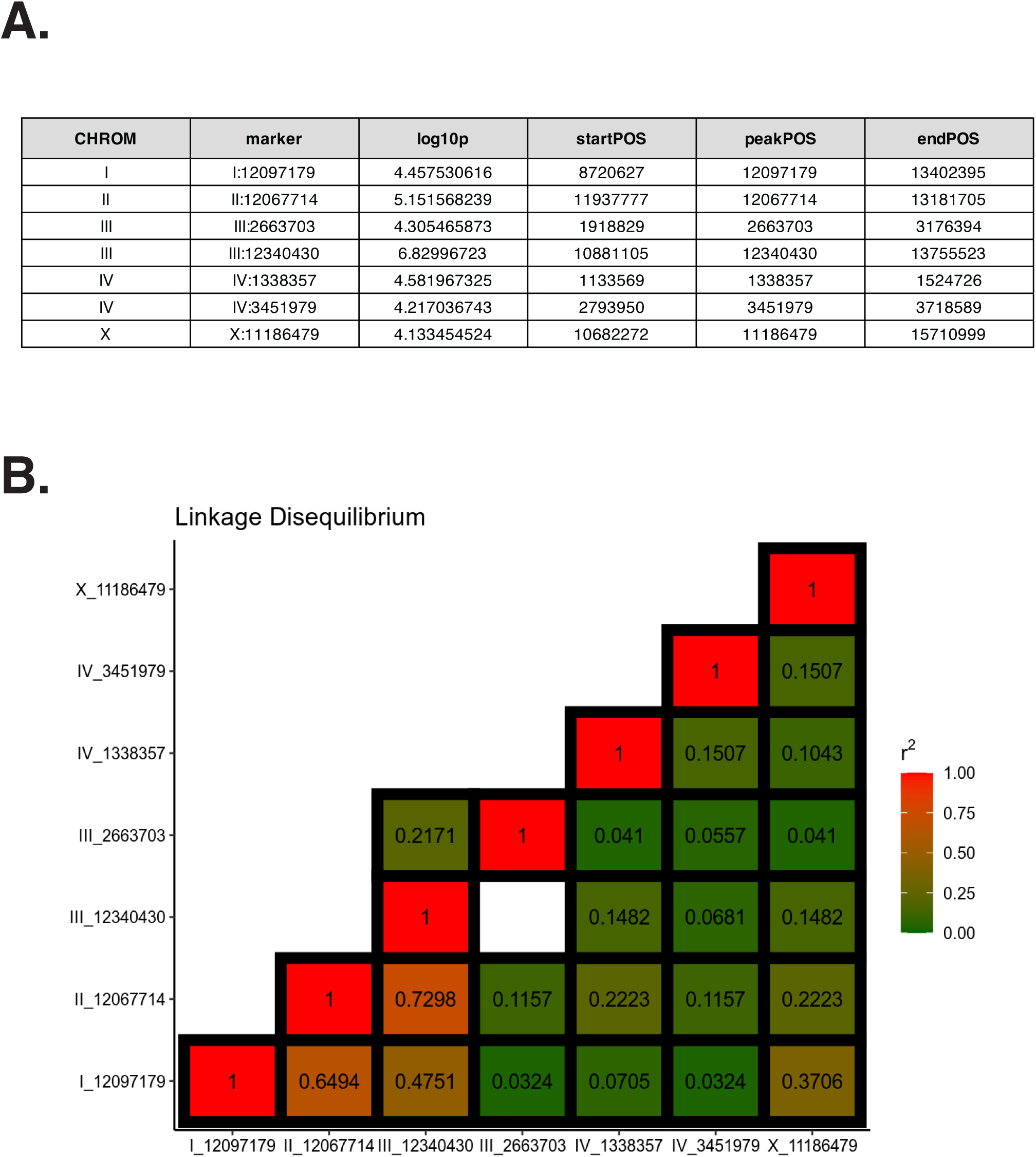
Association mapping of the response against KCN using NemaScan. A. Summary table of QTL peak markers that passed the eigen- decomposition significance (ED) threshold. B. Linkage disequilibrium values between all seven QTL peaks.

**Supplementary Figure 3.**
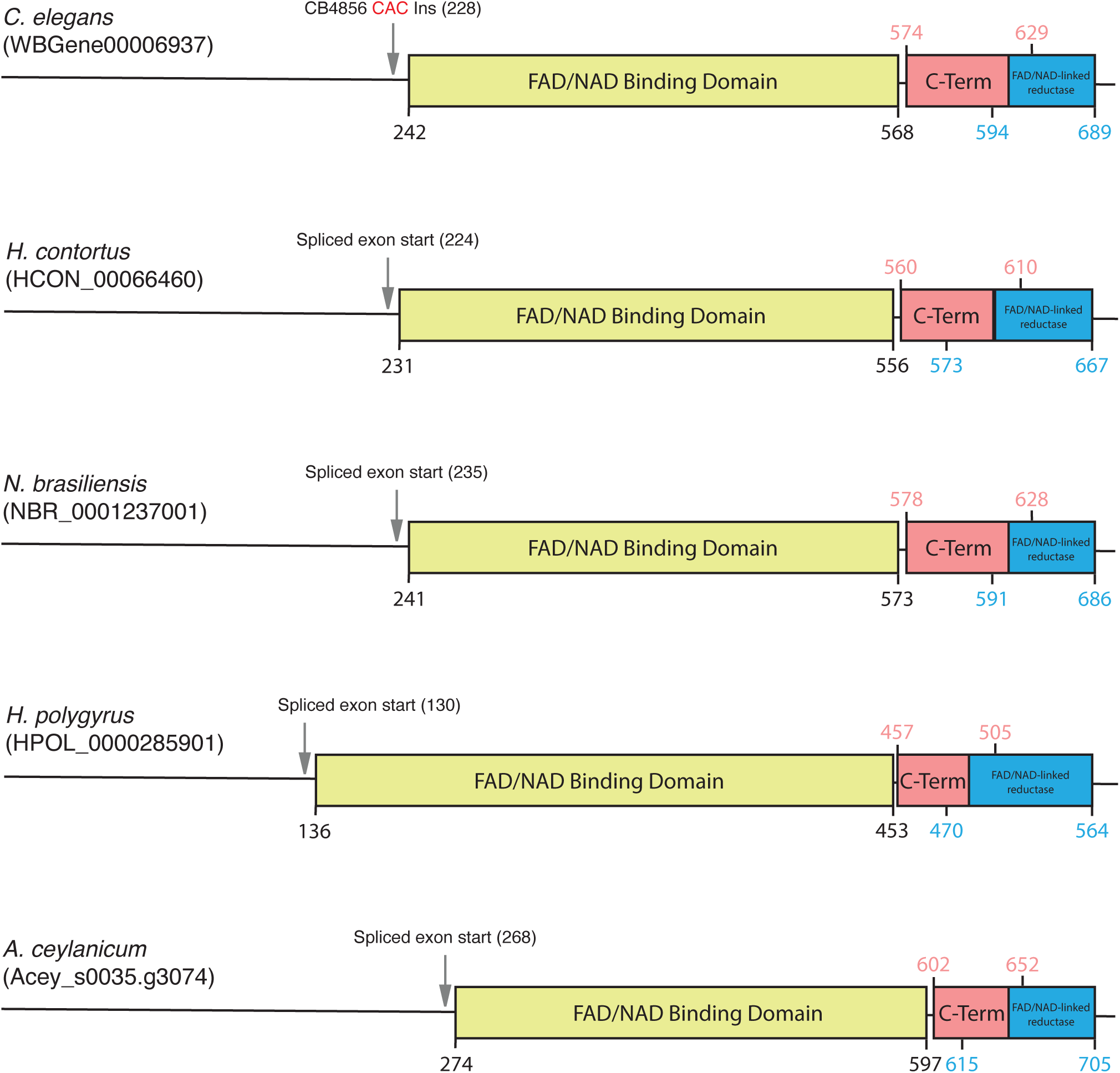
Predicted protein domain architecture in clade V nematode *wah-1* orthologues. Domain predictions as annotated by InterPro. Yellow = FAD/NADH binding domain; pink = C-terminal domain; blue = FAD/NAD-linked reductase domain. Starting position of the spliced isoform highlighted for each orthologue.

**Supplementary Table 1.**
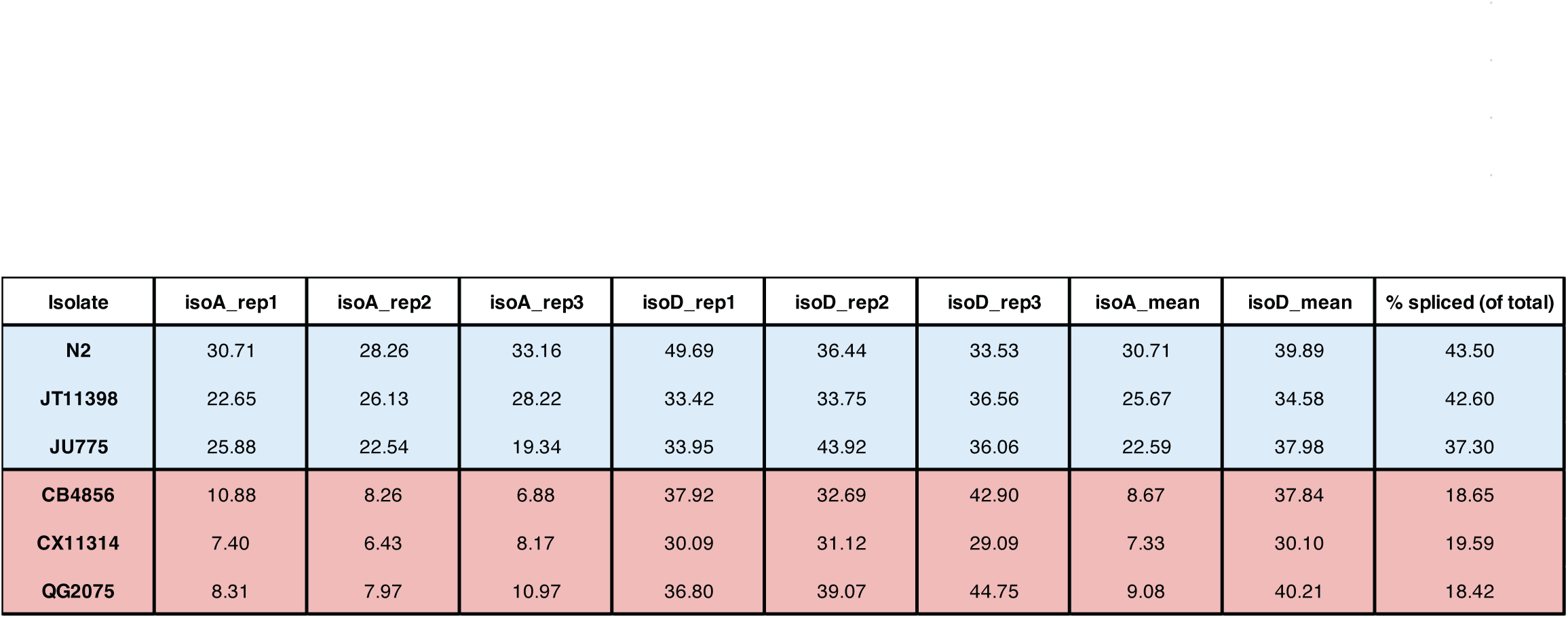
Individual replicates of estimated isoform counts for N2, JT11398, JU775, CB4856, CX11314, and QG2075. Marker used for E3 - III:11994103; marker used for I3 – III:11994104. Percent spliced was calculated as the difference of coverage between E3 and I3 divided by the total coverage over the junction. Percent spliced is calculated with the formula: ((Exon3cov - Intron3cov) / Exon3cov) *100

| Isolate | isoA_rep1 | isoA_rep2 | isoA_rep3 | isoD_rep1 | isoD_rep2 | isoD_rep3 | isoA_mean | isoD_mean | % spliced (of total) |
| --- | --- | --- | --- | --- | --- | --- | --- | --- | --- |
| N2 | 30.71 | 28.26 | 33.16 | 49.69 | 36.44 | 33.53 | 30.71 | 39.89 | 43.50 |
| JT11398 | 22.65 | 26.13 | 28.22 | 33.42 | 33.75 | 36.56 | 25.67 | 34.58 | 42.60 |
| JU775 | 25.88 | 22.54 | 19.34 | 33.95 | 43.92 | 36.06 | 22.59 | 37.98 | 37.30 |
| CB4856 | 10.88 | 8.26 | 6.88 | 37.92 | 32.69 | 42.90 | 8.67 | 37.84 | 18.65 |
| CX11314 | 7.40 | 6.43 | 8.17 | 30.09 | 31.12 | 29.09 | 7.33 | 30.10 | 19.59 |
| QG2075 | 8.31 | 7.97 | 10.97 | 36.80 | 39.07 | 44.75 | 9.08 | 40.21 | 18.42 |

**Supplementary Table 2.**
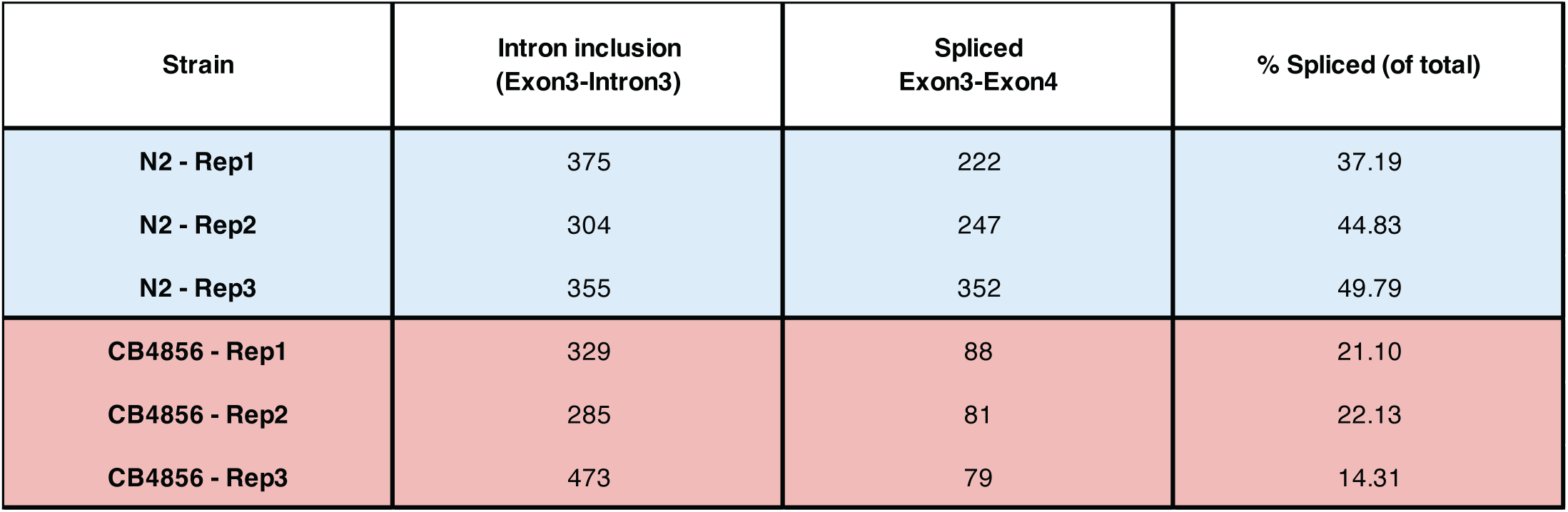
Manual quantification of the RNA-seq reads across the Exon3-Intron3-Exon4 splice junction. Sequencing reads were aligned using HISAT2 and individual splice counts were curated by eye using Integrated Genome Browser.

| Strain | Intron inclusion<br>(Exon3-Intron3) | Spliced<br>Exon3-Exon4 | % Spliced (of total) |
| --- | --- | --- | --- |
| N2 - Rep1 | 375 | 222 | 37.19 |
| N2 - Rep2 | 304 | 247 | 44.83 |
| N2 - Rep3 | 355 | 352 | 49.79 |
| CB4856 - Rep1 | 329 | 88 | 21.10 |
| CB4856 - Rep2 | 285 | 81 | 22.13 |
| CB4856 - Rep3 | 473 | 79 | 14.31 |

**Supplementary Table 3.** Summary of publicly available RNA-seq studies used in this paper. A total of three independent replicates were used for each parasite with the exception of *A. ceylanicum*, that only had one replicate.

| Study | Run | Stage | Organism |
| --- | --- | --- | --- |
| ERP002173 | ERR229528 | Eggs | *Haemonchus contortus* |
| ERP002173 | ERR229529 | Eggs | *Haemonchus contortus* |
| ERP002173 | ERR229530 | Eggs | *Haemonchus contortus* |
| ERP002173 | ERR229531 | L1 | *Haemonchus contortus* |
| ERP002173 | ERR229532 | L1 | *Haemonchus contortus* |
| ERP002173 | ERR229533 | L1 | *Haemonchus contortus* |
| ERP002173 | ERR229534 | L4 | *Haemonchus contortus* |
| ERP002173 | ERR229535 | L4 | *Haemonchus contortus* |
| ERP002173 | ERR229536 | L4 | *Haemonchus contortus* |
| ERP002173 | ERR229537 | Adult female | *Haemonchus contortus* |
| ERP002173 | ERR229538 | Adult female | *Haemonchus contortus* |
| ERP002173 | ERR229539 | Adult female | *Haemonchus contortus* |
| ERP002173 | ERR229540 | Adult male | *Haemonchus contortus* |
| ERP002173 | ERR229541 | Adult male | *Haemonchus contortus* |
| ERP002173 | ERR229542 | Adult male | *Haemonchus contortus* |
| ERP002173 | ERR229546 | Sheathed L3 | *Haemonchus contortus* |
| ERP002173 | ERR229547 | Sheathed L3 | *Haemonchus contortus* |
| ERP002173 | ERR229548 | Sheathed L3 | *Haemonchus contortus* |
| ERP002173 | ERR229549 | Adult female - gut | *Haemonchus contortus* |
| ERP002173 | ERR229550 | Adult female - gut | *Haemonchus contortus* |
| ERP002173 | ERR229551 | Adult female - gut | *Haemonchus contortus* |
| SRP157940 | SRR7693874 | Adult | *Heligmosomoides polygyrus* |
| SRP157940 | SRR7693875 | Adult | *Heligmosomoides polygyrus* |
| SRP157940 | SRR7693876 | Adult | *Heligmosomoides polygyrus* |
| ERP023010 | ERR2023649 | L3 - activated | *Nippostrongylus brasiliensis* |
| ERP023010 | ERR2023650 | L3 - activated | *Nippostrongylus brasiliensis* |
| ERP023010 | ERR2023651 | L3 - activated | *Nippostrongylus brasiliensis* |
| SRP035476 | SRR1124911 | Adult - 19d PI | *Ancylostoma ceylanicum* |

**Supplementary Data 1. Quantitative trait data for 48 CeNDR isolates at 3 hrs post-recovery from KCN incubation**. QT values used for genome-wide association mapping using NemaScan. Data is the mean of at least three biological replicates.

**Supplementary Data 2. List of gene variants found in the QTL on the right arm of chromosome III**. Interval spans from III:10,881,105-13,755,523; Peak marker = III:12,340,430.

